# Effects of immunomodulatory peptides derived from a soil bacterium on cecal microbiota of broilers challenged with *Clostridium perfringens*

**DOI:** 10.1101/2022.03.06.480873

**Authors:** Xiaoying Li, Guiguan Li, Qianning Peng, Yuli Sun, Yong Li, Haitao Li, Jun Ren

## Abstract

*Brevibacillus texasporus* peptide (BT peptide) is immunomodulatory in poultry as a feed additive to substitute antibiotics. In the present study we performed 16S rRNA gene sequencing to compare development of the cecal microbial communities in *Clostridium perfringens*-challenged broilers fed basal diet only or with either BT peptide (48 ppm) or antibiotics mixture (20 ppm bacitracin zinc and 40 ppm colistin sulfate). 240 chicks were randomly assigned into each treatment and a total of 65 cecal samples were collected at the beginning, during seven challenge days and one week post challenge. The composition of microbial communities was clearly distinguishable over time. Treatments with challenge and the antibiotics mixture were associated with increased diversity and with higher relative abundances of *Alistipes* sp. CHKCI003 and *Faecalibacterium* and lower abundance of *Escherichia coli* (all *p* < 0.05). At the termination of the trial, the cecal microbiota in broilers supplemented with BT peptide was dominated by members of Bacteroidaceae. Predicted function analysis reveals significant enrichment of genes involved in ion-coupled transporters and sugar and biotin metabolism in the BT peptide treatment. Taken together, our results suggest that BT peptide and commonly used antibiotics have different influences on modulating the composition of cecal microbiota in broilers.

## 1. Introduction

Probiotics and its functionally valuable products have been shown to be a promising option to modulate the host’s immune system and suggested as an alternative to antibiotics for maintaining animal health (Kanmani et al. 2013). Gram-positive soil bacteria, such as *Streptomyces* and *Amycolatopsis*, could produce antimicrobial peptides (AMPs), and some peptides encrypted in the microbial metaproteome act as natural effectors of the innate immune response (Chen & Lu 2020; Lewies et al. 2019). *Brevibacillus* spp., established in 1996 and formerly known as *Bacillus brevis* cluster (Panda et al. 2014), are rod-shaped Gram-positive bacteria and most of them are strict aerobes. *Brevibacillus* spp. are widely spread in nature and some strains have been applied as probiotics for a long time (Sanders et al. 2003). It is well documented that members of the genus *Brevibacillus* produce antibacterial, antifungal and anti-invertebrate agents and could be a source of diverse enzymes of great biotechnological interests (Yang et al. 2016). One group of these metabolites, *Brevibacillus* AMPs are synthesized through ribosomal or nonribosomal pathway, and the latter involves nonribosomal peptide synthetases (NRPSs). It is interesting to note that the majority of currently-identified *Brevibacillus* AMPs are synthesized by NRPS machinery (Yang & Yousef 2018).

A group of small cationic AMPs, named *Brevibacillus texasporus* peptide (BT peptide), was isolated in 2005 from a soil bacterium *B. laterosporus* in the initial purpose of seeking novel antibiotics (Wu et al. 2005). BT peptide has been found out to be biosynthesized through NRPS and purified to further determine its sequence. Subsequent studies have revealed that the 13-residue BT peptide shares high similarity of amino acid composition with the so-called nonribosomal linear lipopeptides secreted from different *B. laterosporus* strains, including Bogorols A-E and Brevibacillin (Barsby et al. 2006; Yang et al. 2016; Yang & Yousef 2018). Looking into the amino acid sequence alignment, the conserved ornithine in position 3 and lysines in positions 7 and 10 make this family of peptides cationic at physiological pH. Several evidences suggest that BT peptide has immunomodulatory properties that prime innate immunity and enhance leukocyte bactericidal activity to combat pathogenic infection when provided as a feed additive for broilers (Kogut et al. 2007; Kogut et al. 2012; Kogut et al. 2010). However, there are no studies, to our knowledge, examining the effects of BT peptide on gut microbiota composition of the host so far.

Necrotic enteritis (NE) is a gastrointestinal disease of broiler chickens caused by *Clostridium perfringens* strains (Lacey et al. 2018). Subclinical NE is characterized by poor performance with no clinical sign of the disease (Wang et al. 2017). In this study, we applied a NE challenge model with broiler chickens to investigate the role of BT peptide in gut bacterial community structure shifts and compare it with the antibiotics commonly used for eradication of the pathogenic bacteria. High-throughput sequencing of 16S rRNA genes was performed to follow the response of the cecal microbial communities to *C. perfringens* challenge.

## 2. Materials and Methods

### 2.1. Animals

All procedures involving animals were carried out according to experimental protocols approved by the Animal Ethics Committee of COFCO Nutrition and Health Research Institute (permit number: AR-15-0701). A total of 240 one-day-old male Arbor Acres broiler chickens were obtained and transferred from a commercial hatchery (Huadu Broiler Breeding, Beijing, China) and randomly distributed into three treatments, each group with 80 birds housed in individual cage compartments within a climate-controlled room. Each compartment had its own feeder, drinker, and brooding lamp for warmth, and plastic mesh was used for the cage floor. The ambient relative humidity inside the room was around 50%, and the environmental temperature was 32°C at placement being reduced with age to provide comfort throughout the study. Treatments were as follows: basal starter diet (coccidiostat- and antibiotic-free, referred to as CP hereafter), 20 ppm bacitracin zinc + 40 ppm colistin sulfate (AB), and 48 ppm BT peptide (BT), and the feeding period lasted 18 days. Feed and water were supplied under *ad libitum* strategy and the chickens were maintained on a 23-hour lighting program. The corn and soybean meal-based diets were formulated to meet NRC (1994) requirements. Composition of the diet and nutrient levels are presented in Table 1.

**Table 1.** Ingredients and nutrient composition of the diet (as-fed basis)

|  | Value |
| --- | --- |
| **Ingredient, %** |  |
| Corn | 57.2 |
| Soybean meal, 46% CP | 34.6 |
| Soybean oil | 3.0 |
| Dicalcium phosphate | 1.7 |
| Limestone | 1.4 |
| 2% Premix<sup>a</sup> | 2.0 |
| Antioxidant | 0.1 |
| **Nutrient composition** |  |
| Metabolizable energy, MJ/kg | 12.33 |
| Crude protein, % | 21.00 |
| Lys, % | 1.20 |
| Met, % | 0.50 |
| Ca, % | 1.05 |
| Available P, % | 0.45 |

<sup>a</sup>The premix supplies the following per kg of complete feed: vitamin A, 12,500 IU; vitamin D3, 2,500 IU; vitamin K3, 2.65 mg; vitamin B1, 2 mg; vitamin B2, 6 mg; vitamin B12, 0.025 mg; vitamin E, 30 IU; biotin, 0.0325 mg; folic acid, 1.25 mg; pantothenic acid, 12 mg; niacin, 50 mg; copper, 8 mg; zinc, 75 mg; iron, 80 mg; manganese, 100 mg; selenium, 0.15 mg; iodine, 0.35 mg.

*C. perfringens* challenge was adapted from published methods (Liu et al. 2010) with some modifications. Briefly, a chicken *C. perfringens* type A field strain was obtained from the China Veterinary Culture Collection Center (Beijing, China) and cultured anaerobically in cooked meat medium (Aobox, Beijing, China) overnight at 37°C. At 4 to 10 days posthatch, all the birds were orally gavaged once per day at approximately the same time of morning with the actively growing cultures of this pathogenic *C. perfringens* (2.0 × 10^8^ cfu/ml, 1.0 ml/bird). Plate count of viable *C. perfringens* was done by performing serial dilution of the culture in sterile saline and plated on sulfite-polymyxin-sulfadiazine agar (Aobox). The plates were incubated anaerobically at 37°C for 18–24 hours and colonies typical for *C. perfringens* were counted.

### 2.2. Sample collection

Samples were collected at afternoon for each sampling time. At 4 (start of challenge, referred to as T1 hereafter), 11 (1 d post-challenge, T2), and 17 (7 d post-challenge, T3) days posthatch, 6□8 birds from each treatment were randomly selected and humanely euthanized by cervical dislocation. Thus, 9 sampling groups were generated, namely CP1, CP2, CP3, AB1, AB2, AB3 and BT1, BT2, BT3 according to treatment and sampling sequence. Mucoid gut contents and sparse occurrence of focal necrosis in the small intestine with no obvious mortality implied a successful challenge. From each chicken, the cecum was removed and the cecal luminal contents were collected in a sterilized tube. The samples were flash frozen and stored at −80°C until further analysis.

### 2.3. DNA isolation and sequencing

Bacterial genomic DNA of each sample was extracted using the QIAamp Fast DNA Stool Mini Kit (Qiagen, Hilden, Germany) following the manufacturer’s instructions. DNA concentration and purity were monitored by agarose electrophoresis and determined by UV absorption analysis using NanoDrop spectrophotometer (Thermo Fisher Scientific, Waltham, USA). The V4 hypervariable region of the 16S rRNA bacterial gene (515-806) was amplified using specific primers with the barcodes with Phusion High-Fidelity PCR Master Mix (New England Biolabs, Ipswich, USA). PCR amplicons from each sample were pooled in equimolar amounts and purified using QIAquick Gel Extraction Kit (Qiagen). Sequencing libraries were generated using TruSeq DNA PCR-Free Sample Prep Kit (Illumina, San Diego, USA) and assessed by Qubit 2.0 Fluorometer (Thermo Fisher Scientific) and Agilent 2100 Bioanalyzer system according to the standard protocols. The final library was paired-end sequenced at 2 × 250 bp on the Illumina HiSeq 2500 platform. The raw sequence data are available at the NCBI Sequence Read Archive (SRA) database (Accession Number: SRP144400).

### 2.4. Sequence analyses

Raw sequences were demultiplexed and quality-filtered using QIIME (version 1.7.0, http://qiime.org) (Caporaso et al. 2010) to eliminate all low quality sequence reads under specific filtering conditions (Bokulich et al. 2013). The resulting trimmed sequences were filtered to remove singleton reads and then grouped into operational taxonomic units (OTUs) with 97% identity threshold using the USEARCH software based on the UPARSE algorithm (version 7.1, http://drive5.com/uparse) (Edgar 2013) while chimeric sequences were identified and removed using UCHIME (Edgar et al. 2011). The taxonomy of OTU-representative sequences was analyzed by the RDP Classifier (version 2.2, http://rdp.cme.msu.edu) (Wang et al. 2007) against the Silva (SSU128) 16S rRNA database using confidence threshold of 70%. The resulting OTU table was used to determine taxonomic relative abundances of each sample.

### 2.5. Statistical analyses

Data analyses were performed with SPSS Statistics software (version 24.0, IBM Corporation, Armonk, USA) and R (version 3.3.1, http://www.r-project.org). Non-parametric statistical methods were used to analyze the data sets when they don’t follow the normal distribution. Differences in α-diversity for the cecal microbiota of broilers were tested using Kruskal-Wallis H test with Dunn’s post hoc test and Bonferroni correction for multiple comparisons. Differences in microbiota composition as assessed by β-diversity metrics were tested using analysis of similarities (ANOSIM) or permutational multivariate analysis of variance (PERMANOVA, 999 permutations) at OTU level in the R vegan package (version 2.4-4, http://CRAN.R-project.org/package=vegan). Differences in abundance of taxa were tested using Kruskal-Wallis H test with *p*-values adjusted (*p_adj_*) for multiple testing by the FDR procedure. Linear discriminant analysis effect size (LEfSe) (Segata et al. 2011) was performed to identify the Kyoto Encyclopedia of Genes and Genomes (KEGG) pathways differentially represented among groups.

## 3. Results

We reproduced the subclinical necrotic enteritis disease in newborn broiler chickens by treatment of 7-consecutive-day *C. perfringens* challenge and followed longitudinally the development in the composition of cecal microbiota (Fig. 1a). Cecal digesta samples were processed and sequenced to generate 3,535,329 clean reads using the 16S rRNA gene’s V4 hypervariable region (Table S1). After OTU clustering and rarefying to 37,970 reads per sample, 1,635 OTUs were retained for downstream analyses.

**Figure 1.**
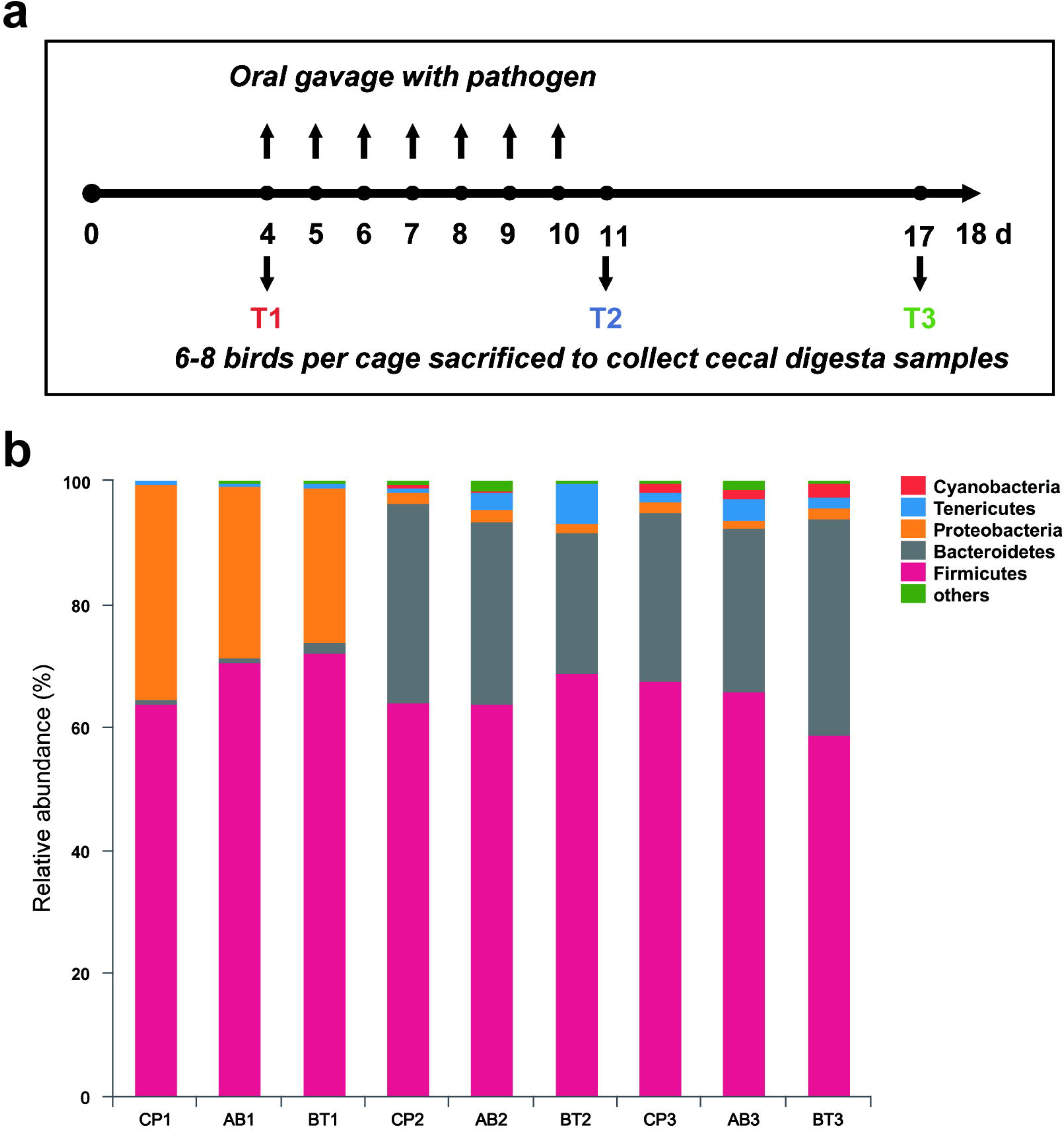
Microbial community composition of the cecum of broiler chickens challenged with *C. perfringens*. (a) Scheme of sampling strategy. T1-T3 represent the three sampling time points in this study, generating a total of nine groups for the following analyses. (b) Bar plots show the mean values of relative abundance for the most abundant phyla in the nine sampling groups. Only phyla present in at least 1% of the samples are shown separately.

### 3.1. Impact of treatments and *C. perfringens* challenge on composition profile and alpha diversity of the bacterial community

The cecal microbiota profiles of challenged broilers at phylum level are shown in Fig. 1b. It can be observed that the phylum Proteobacteria was highly abundant in the T1 samples of all the three treatments but declined to a small portion in the T2 and T3 samples, while replaced by another phylum Bacteroidetes in these samples.

Next, alpha diversity was calculated to describe the within-sample bacterial richness and diversity using observed OTUs and Shannon index. When examining within different time points, we found that AB1 group had a significantly higher number of observed OTUs than CP1 in T1 sampling time (*p* = 0.037, Fig. 2a). We also compared alpha diversity indices in different treatments to evaluate the impact of challenge. It is shown that both bacterial community richness (*p* = 0.018, Fig. 2b) and diversity (*p* = 0.002, Fig. 2c) were notably increased over time in CP treatment. Moreover, Shannon index in AB treatment also exhibited a gradual increase during the entire course of treatment (*p* = 0.029, Fig. 2d).

**Figure 2.**
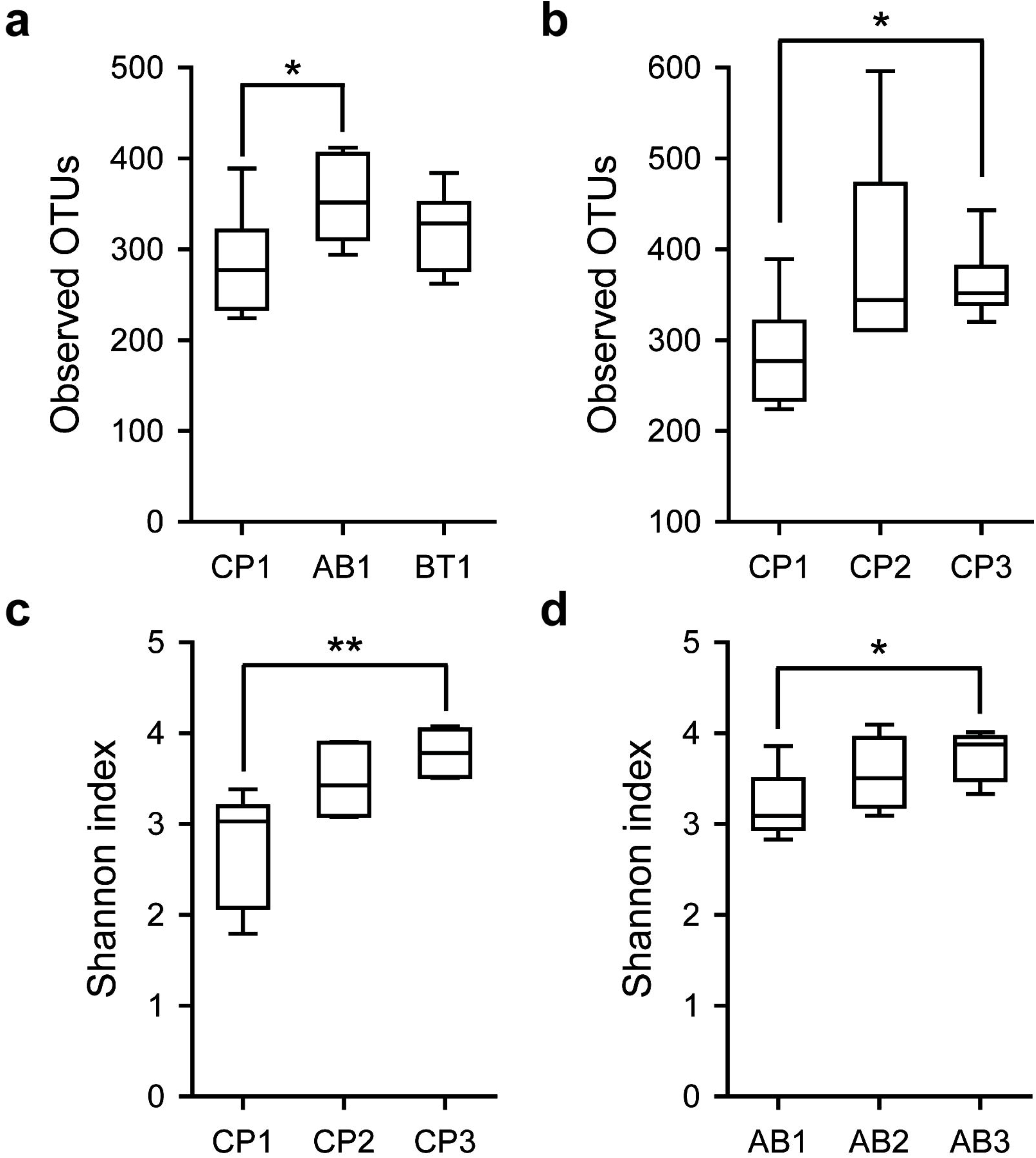
Differences in bacterial community richness and diversity between treatments and sampling times. The bacterial abundance was represented by the numbers of OTUs and compared showing impact of treatment in T1 sampling time (a) or impact of challenge in CP treatment (b). Shannon index was used to estimate the diversity of the cecal microbiota of the challenged birds in CP (c) or AB (d) treatment. *, *p* < 0.05; **, *p* < 0.01.

### 3.2. Cecal bacterial community composition changed over time in challenged broilers

Bacterial community composition was significantly different between sampling groups formed by treatments and time when measured by ANOSIM distances (weighted UniFrac, *r* = 0.514, *p* = 0.001). These differences were recapitulated in two dimensional non-metric multidimensional scaling analysis (NMDS) of the data based on weighted UniFrac similarity distance, which reveals a clear cut grouping structure of taxonomic composition over sampling time (Fig. 3a). We then investigated the cecal microbiota at the genus level to identify the 30 most abundant taxa (Fig. 3b). The heat map graphically shows that the cecal bacterial community in the nine sampling groups varied, especially observed in the course of time, which supports the ANOSIM and NDMS analyses.

**Figure 3.**
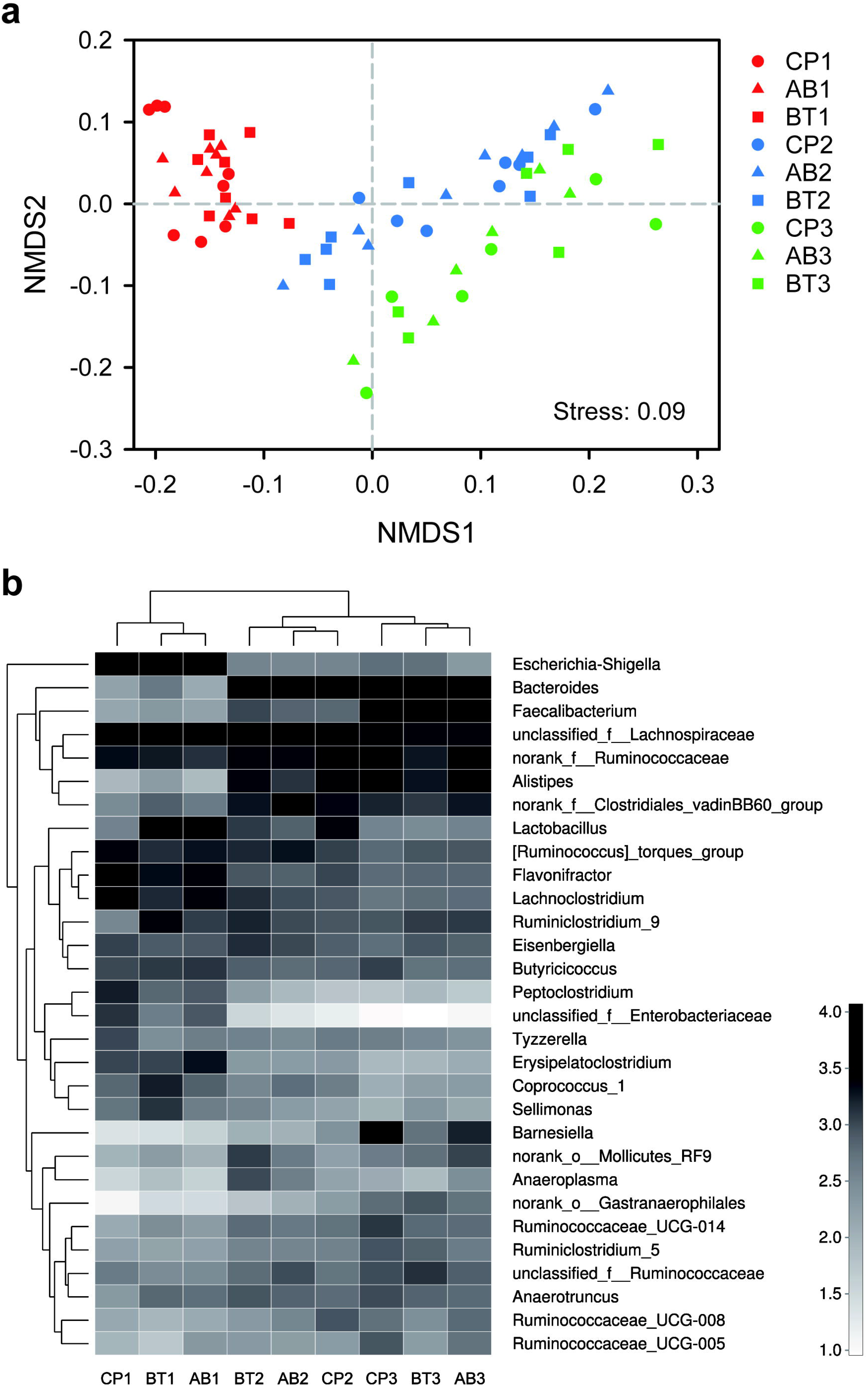
Changes in cecal bacterial community composition over time in the broilers challenged with *C. perfringens*. (a) NMDS ordination based on weighted UniFrac similarity at T1 (red), T2 (blue) and T3 (green) sampling times. (b) Heatmap based on the relative abundance of thirty most abundant genera in the nine sampling groups. Clustering of groups (top) and taxa (left) is defined by the average pairwise Euclidean distance.

To single out the contribution of individual OTUs to the dissimilarity among samples, we assessed taxonomic changes that differed significantly over time in each treatment (Fig. 4). Surprisingly, the abundance of OTU284 (*Escherichia coli*) substantially decreased above 20-fold from T1 to T2 and T3 sampling times in both CP (*p_adj_* = 0.036) and AB (*p_adj_* = 0.049) treatments. This trend coincided with the increases in the abundances of OTU1111 (*Alistipes* sp. CHKCI003) and OTU1175 (*Faecalibacterium*) over the time course of the study. In CP treatment, OTU1092 (Ruminococcaceae) and OTU1052 (*Lactobacillus*) were observed to increase from T1 to T2 sampling times and then decline from T2 to T3 sampling times. In contrast, the abundance of OTU1052 decreased over time in AB treatment groups (*p_adj_* = 0.045), and OTU304 (*Lachnoclostridium*) showed a similar pattern of reduced abundance (*p_adj_* = 0.043).

**Figure 4.**
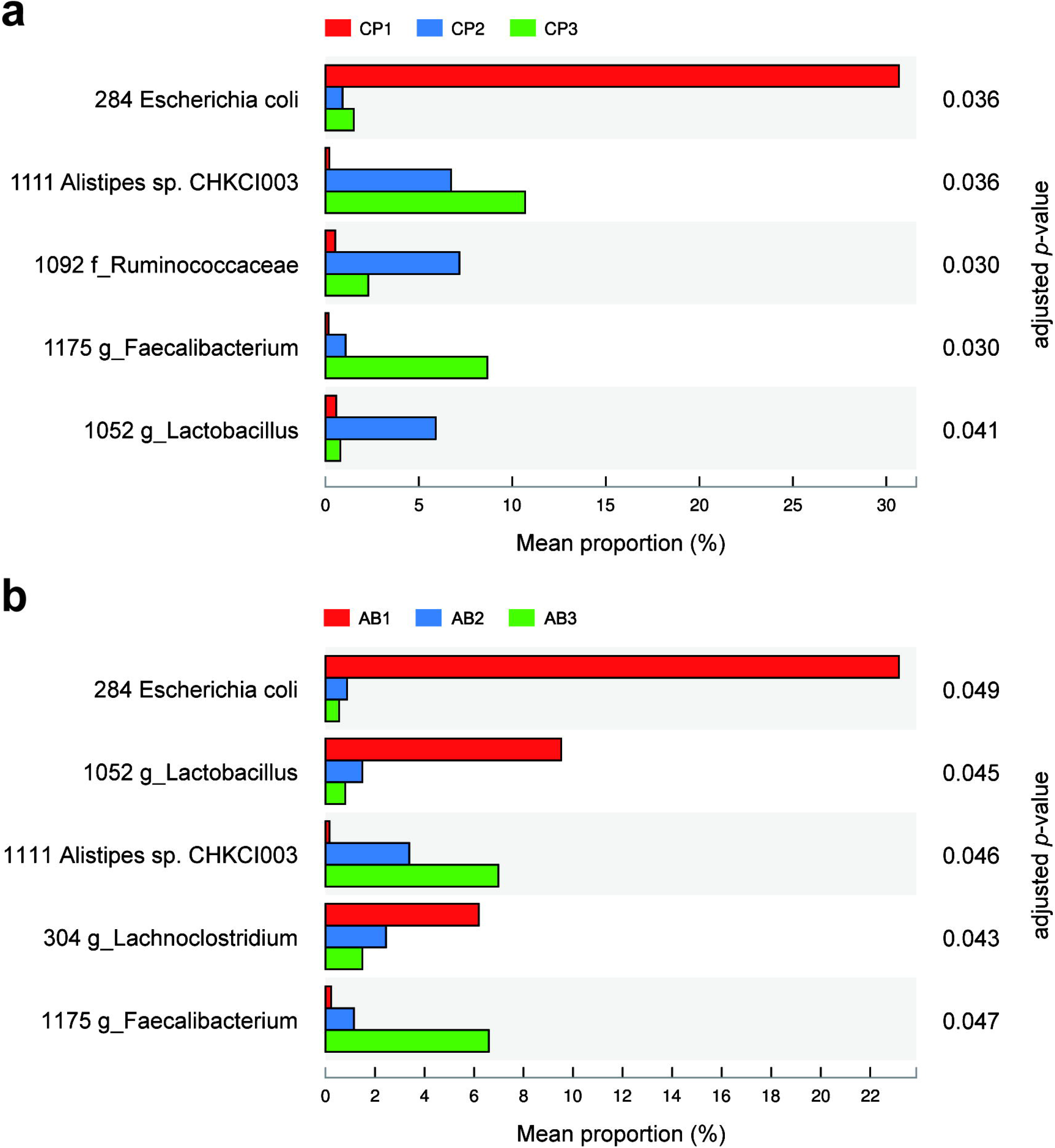
Bar plots identifying significant differences between mean proportions of bacterial taxa in CP (a) and AB (b) treatments over time (*p_adj_* < 0.05). Only OTUs at least 5% abundant in at least one group are included.

### 3.3. Shift in bacterial communities caused by different treatments at T3 sampling time

To assess the impact of different treatments on the cecal microbial community of challenged broiler chickens, Bray-Curtis similarity-based principal coordinate analysis (PCoA) was performed on sequencing data of T3 samples and the first three coordinates are shown in Fig. 5a (representing 63% of the total variance). This analysis indicates a clear division between CP3 and BT3 groups, and the results of PERMANOVA analysis on OTU level also reveal a modest global variation in bacterial composition among different treatment groups (Bray-Curtis, *r*^2^ = 0.222, *p* = 0.013). We then analyzed the data using ternary plots to show the relative distribution of OTUs among T3 samples (Fig. 5b). Each OTU was categorized as group representative OTUs based on whether there was a defined (10% or above) increase on its relative abundance in this treatment group compared with the other two groups. Our results suggest that CP3 and AB3 had a high relative abundance of Ruminococcaceae, whereas BT3 harbored a high relative abundance of Bacteroidaceae, which was solely contributed by OTU810 (*Bacteroides dorei*; see Table S2 for a complete list), and a less relative abundance of Ruminococcaceae. OTUs from the families Rikenellaceae and Porphyromonadaceae were mostly associated with the CP3 bacterial community, whereas Clostridiales_vadinBB60_group was associated with the AB3 community.

**Figure 5.**
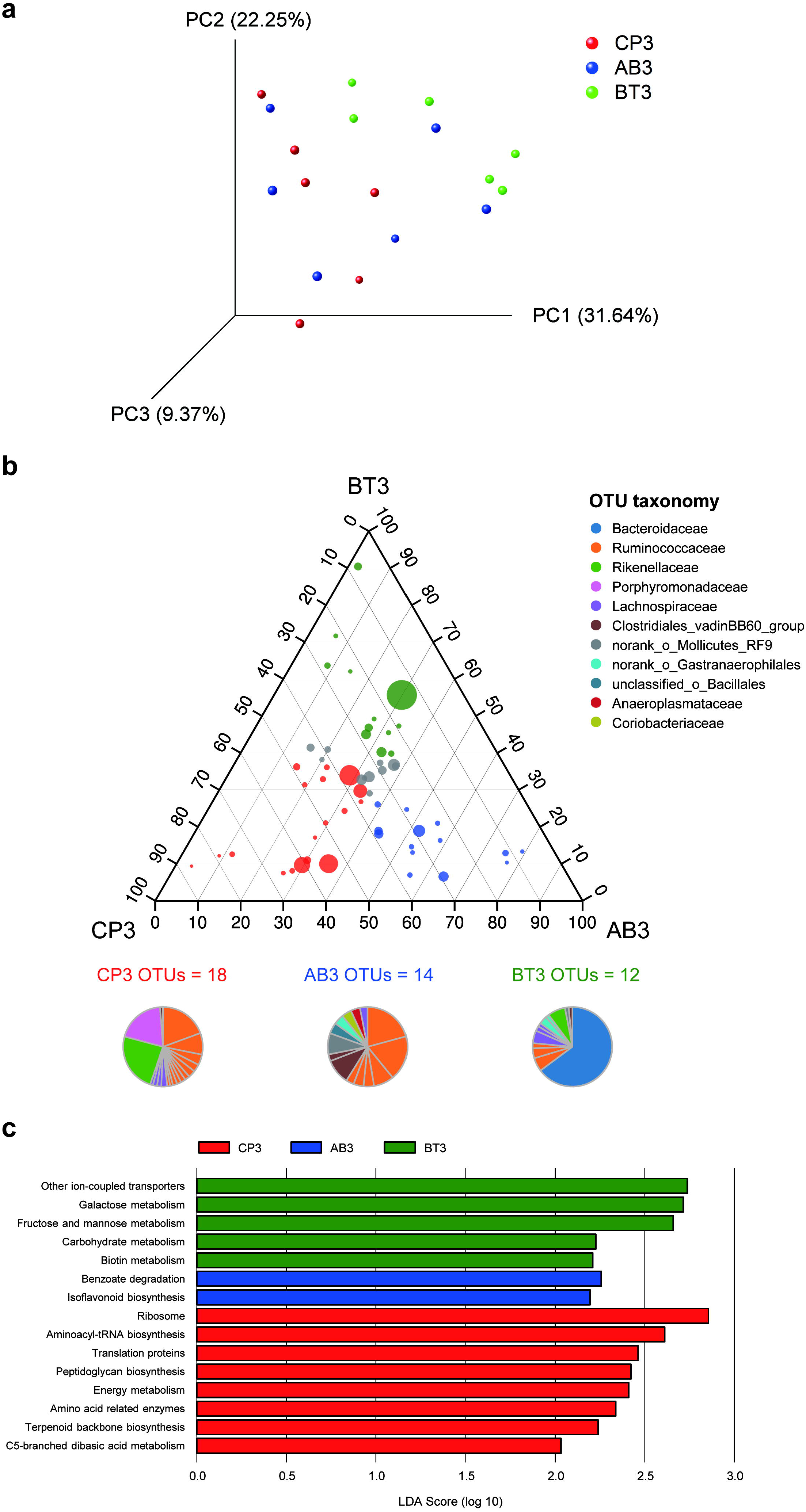
Taxonomic differences and predicted function of the cecal microbiota in the differently treated broilers challenged with *C. perfringens* at T3 sampling time. (a) PCoA plot based on Bray-Cutis distance at OTU level illustrating the distribution of microbiota by treatment group. The percentage of variation explained by the principal coordinates is indicated on the axes. (b) Ternary plot representing the relative occurrence of individual OTUs (circles) in T3 samples. The size of the circles is proportional to the mean abundance of each OTU, and the position of the circles is determined by the contribution of the indicated treatments to the total relative abundance. Red, blue and green circles mark OTUs at least 10% more enriched in CP3, AB3 and BT3 than the other two groups respectively, and their relative abundance is shown as pie charts and colored by family-level taxonomic classification. Only OTUs at least 0.5% abundant in at least one treatment are shown. (c) KEGG level 3 pathways significantly differentiated among treatments at T3 sampling time identified by LEfSe with cutoff of linear discriminant analysis (LDA) > 2.

To study the functional alterations of the cecal microbiota, we used the Phylogenetic Investigation of Communities by Reconstruction of Unobserved States (PICRUSt) (Langille et al. 2013) to predict the functional composition profiles from the 16S rRNA sequencing data in all treatment groups at T3 sampling time. It is found that multiple KEGG level 3 categories were enriched in different treatments (Fig. 5c). The pathways overrepresented in the CP3 group highlighted protein translation (ribosome, aminoacyl-tRNA biosynthesis, translation proteins, and amino acid related enzymes) and four metabolism pathways (peptidoglycan biosynthesis, energy metabolism, terpenoid backbone biosynthesis, and C5-branched dibasic acid metabolism). In contrast, the gut microbiota of the BT3 group was characterized by enrichment of other ion-coupled transporters and metabolism of sugars (galactose, fructose and mannose, and carbohydrate) as well as biotin, whereas the gut microbiota of the AB3 group was characterized by overrepresentation of benzoate degradation and isoflavonoid biosynthesis.

## 4. Discussion

The purpose of this study was to define the microbiome of broilers challenged with *C. perfringens*, and to evaluate the impact of BT peptide on the broiler microbiome in comparison with the antibiotics mixture.

Necrotic enteritis caused by *C. perfringens* in broilers is a contagious disease of economic importance worldwide (Yitbarek et al. 2012). The disease has two forms, clinical and subclinical, and most of the economic losses are related to the subclinical form and the high cost of preventing the disease with antibiotics (Shojadoost et al. 2012). In recent years, countries and regions where in-feed antimicrobials or antibiotic growth promoters have been banned have experienced an increase in disease outbreaks in broiler flocks (Van Immerseel et al. 2004). Thus, it has spurred interest in investigating how it can be prevented by countermeasures other than the use of antibiotics.

BT peptide has been proposed as a feed additive for poultry to achieve effective control of food safety pathogens. It is evident that feeding the BT peptide-supplemented diet in neonatal broiler chickens induced the up-regulation of the innate immune response, reduced pathogenic bacterial colonization of the intestine and primed the cecal tissue for increased immune gene expression in response to *Salmonella enterica* serovar Enteritidis infection (Kogut et al. 2007; Kogut et al. 2010). Researchers have realized that the large numbers of microorganisms residing in the gastrointestinal tract have a highly coevolved relationship with the host’s immune system (Hooper et al. 2012), supported by the fact that disturbances in the bacterial microbiota result in dysregulation of immune cells (Round & Mazmanian 2009). Particularly, it is well understood that the interaction between the gut immune system and commensal microbes in chickens starts immediately after hatching (Crhanova et al. 2011). Since BT peptide is not absorbed in the intestine (Kogut et al. 2012), we considered the effect of its supplementation on diversity and function of the avian gut microbiota.

As described in results, the counteracting relationship of Proteobacteria and Bacteroidetes observed between T1 and T2/T3 can be partially explained by the high existence of the genus *Escherichia-Shigella* in the first week of age and *Bacteroides* thereafter, which shares strong similarity with findings of previous research (Johnson et al. 2018; Waite & Taylor 2014). A core bacterial microbiota of broiler cecum was proposed comprising *Escherichia-Shigella* at 0 and 7 days of age and *Bacteroides*, Erysipelotrichaceae, *Faecalibacterium*, Lachnospiraceae, *Oscillospira*, *Rikenella*, *Ruminococcus*, *Streptococcus* and *Lactobacillus* spp. after 14 days of age (Johnson et al. 2018; Stanley et al. 2014). Temporal changes in the chicken cecal microbiota suggest that taxonomic richness and diversity typically increase from day of hatch to maturity (Oakley & Kogut 2016), as revealed by our alpha diversity analysis.

The pathogenic strain that was used for challenging broiler chickens belongs to the genus *Clostridium*, which could be a predominant bacterium in the cecum depending on the age of the bird (Stanley et al. 2012). However, we did not detect a marked colonization of the cecum with *C. perfringens* throughout the sampling time. One of the possible reasons is that the gross lesions caused by *C. perfringens* infection usually happen in the small intestine including jejunum and ileum (Shojadoost et al. 2012), thus in the cecal mucosa may not be the primary site for the pathogen to colonize. We also notice that a low relative abundance of *Clostridium* in 16S rRNA sequencing data and low cecal counts of *C. perfringens* were obtained in a study on broilers at 0-36 days of age (Ranjitkar et al. 2016), implying a difficult colonization of the cecal environment for *C. perfringens* in the early stage of broiler chickens.

Analysis of the microbiome at 7 d post-challenge shows that the most abundant organisms were discriminating among different treatments. For challenged broilers fed with BT peptides, the dominant taxa were *Bacteroides dorei* and Ruminococcaceae. *Bacteroides* colonization of the gastrointestinal tract at early stage of life is important given their role in the digestion of complex carbohydrates to fermentation products which are beneficial to hosts (Davis-Richardson et al. 2014; Martens et al. 2008). Notably, our PICRUSt results also reveal that predicted genes from the metagenome related to multiple carbohydrate metabolic pathways were significantly increased in BT treatment. *Bacteroides dorei*, particularly, has been found to exhibit strong association with the development of autoimmunity in type 1 diabetes in a Finnish cohort of children (Davis-Richardson et al. 2014). The structurally and functionally distinct lipopolysaccharide of *Bacteroides dorei* is assumed to play an important role in immune regulation in human individuals (Vatanen et al. 2016). This study lacks sufficient data to define the effect of high *Bacteroides* abundance on broiler immune response to *C. perfringens* pathogen, and further studies are needed to investigate this potential link.

Various bacteria belonging to Ruminococcaceae were most abundant in both CP and AB treatments at 7 d post-challenge. Members of the family Ruminococcaceae including *Faecalibacterium* are known for their ability to decompose plant material and convert into butyrate and other short-chain fatty acids that can be absorbed and used for energy by the host. Significant enrichment of pathways regulating energy metabolism in CP treatment also demonstrates a heightened demand for energy by the cells of the intestine in this condition. Furthermore, we identified two *Bacteroidetes* OTUs, OTU1111 (*Alistipes* sp. CHKCI003, a species from the family of Rikenellaceae) and OTU377 (*Barnesiella*, a genus from the family of Porphyromonadacea*e*), which were more enriched in CP treatment. These two bacteria have been reported to contribute to host immune development. The appropriate proportion of *Alistipes* is suspected to augment intestinal immune maturation (Chung et al. 2012), and administration of the commensal *Barnesiella* could eradicate the infection of vancomycin-resistant *Enterococcus* (Ubeda et al. 2013), suggesting the microbiota as a required component of the effector response of the host (Belkaid & Hand 2014). Nevertheless, we need more research to unravel the underlying mechanism of this bacterial community development.

Collectively, our results show that feeding *C. perfringens*-challenged broilers with BT peptide or the antibiotics causes a different outcome on the composition of microbial community in cecum. Additionally, analysis of the predicted metagenomes indicates functions involved with improved sugar metabolism when supplemented with BT peptide. Future studies should include a more explicit investigation of the host-microbe interaction to depict a detailed picture of the immunomodulatory activity of BT peptide from the perspective of gut microbiota, which may benefit BT peptide as a novel antibiotics alternative for a range of livestock species.

## Acknowledgements

This work was supported by the Beijing Municipal Science & Technology Program under Grant Z141100002614014; the National Key R&D Program of China under Grant 2018YFD0500600; and the Basic Research Funds of COFCO Nutrition and Health Research Institute. We appreciate Dr. Yiwei Jiang (My Galaxy LLC, Ft. Worth, USA) for providing the BT peptide material for this study. We would like to thank Dr. Yuming Guo (China Agricultural University, Beijing, China) for technical assistance with the challenge experiment and Dr. Shuangli Mi (Beijing Institute of Genomics, Chinese Academy of Sciences, Beijing, China) for assistance with sample sequencing during the study.

