## Supplementary material for "Effects of immunomodulatory peptides derived from a soil bacterium on cecal microbiota of broilers challenged with *Clostridium perfringens*": Table S1

**Table S1. OTU table summary**

| **Sample ID** | **Reads** | **OTUs** | **Good’s** | **ACE** | **Chao1** | **Shannon** | **Simpson** |
| --- | --- | --- | --- | --- | --- | --- | --- |
| CP1_1 | 40019 | 333 | 99.78% | 422 | 430 | 3.2058 | 0.1212 |
| CP1_2 | 51372 | 272 | 99.78% | 458 | 411 | 3.3808 | 0.0652 |
| CP1_3 | 63244 | 283 | 99.79% | 426 | 371 | 2.2854 | 0.2434 |
| CP1_4 | 48003 | 233 | 99.82% | 315 | 322 | 1.7919 | 0.3971 |
| CP1_5 | 56013 | 282 | 99.76% | 518 | 435 | 2.9842 | 0.1237 |
| CP1_6 | 51012 | 224 | 99.87% | 266 | 283 | 2.0049 | 0.3481 |
| CP1_7 | 61880 | 238 | 99.83% | 318 | 310 | 3.1601 | 0.0767 |
| CP1_8 | 61892 | 389 | 99.67% | 625 | 541 | 3.0734 | 0.1162 |
| AB1_1 | 51764 | 351 | 99.73% | 537 | 479 | 2.9797 | 0.1572 |
| AB1_2 | 60528 | 412 | 99.71% | 513 | 540 | 3.8581 | 0.0434 |
| AB1_3 | 64532 | 352 | 99.69% | 610 | 491 | 3.0353 | 0.1420 |
| AB1_4 | 44994 | 294 | 99.79% | 448 | 399 | 3.3780 | 0.0747 |
| AB1_5 | 57817 | 412 | 99.56% | 827 | 629 | 3.1393 | 0.1014 |
| AB1_6 | 51570 | 332 | 99.75% | 431 | 416 | 2.8291 | 0.1294 |
| AB1_7 | 48737 | 384 | 99.67% | 626 | 552 | 3.5324 | 0.0549 |
| AB1_8 | 48228 | 304 | 99.79% | 439 | 402 | 2.9303 | 0.1568 |
| BT1_1 | 44931 | 345 | 99.73% | 472 | 479 | 2.7157 | 0.2154 |
| BT1_2 | 46790 | 346 | 99.73% | 546 | 456 | 3.0883 | 0.1439 |
| BT1_3 | 52389 | 277 | 99.81% | 352 | 350 | 2.8514 | 0.1436 |
| BT1_4 | 50693 | 277 | 99.79% | 371 | 394 | 3.4186 | 0.0564 |
| BT1_5 | 61725 | 262 | 99.82% | 380 | 353 | 3.3436 | 0.0775 |
| BT1_6 | 52148 | 312 | 99.80% | 395 | 426 | 3.3185 | 0.0871 |
| BT1_7 | 48238 | 384 | 99.75% | 477 | 511 | 3.8531 | 0.0464 |
| BT1_8 | 57252 | 353 | 99.74% | 467 | 508 | 3.2927 | 0.1167 |
| CP2_1 | 56674 | 596 | 99.56% | 756 | 742 | 3.0881 | 0.2175 |
| CP2_3 | 53157 | 311 | 99.81% | 380 | 402 | 3.3280 | 0.1071 |
| CP2_4 | 47897 | 311 | 99.79% | 403 | 402 | 3.0800 | 0.1277 |
| CP2_5 | 65501 | 344 | 99.74% | 549 | 527 | 3.9038 | 0.0445 |
| CP2_6 | 56235 | 472 | 99.62% | 648 | 632 | 3.8960 | 0.0474 |
| CP2_7 | 59356 | 338 | 99.81% | 411 | 411 | 3.6371 | 0.0869 |
| CP2_8 | 61484 | 360 | 99.77% | 452 | 451 | 3.4270 | 0.1365 |
| AB2_1 | 46058 | 699 | 99.59% | 822 | 874 | 3.5132 | 0.1799 |
| AB2_2 | 44661 | 320 | 99.80% | 453 | 409 | 3.8736 | 0.0421 |
| AB2_3 | 47646 | 365 | 99.76% | 462 | 491 | 3.9740 | 0.0350 |
| AB2_4 | 56402 | 324 | 99.77% | 421 | 435 | 3.4906 | 0.0803 |
| AB2_5 | 40456 | 494 | 99.72% | 581 | 613 | 4.0934 | 0.0528 |
| AB2_6 | 61127 | 340 | 99.76% | 519 | 498 | 3.0898 | 0.2022 |
| AB2_7 | 56965 | 325 | 99.81% | 400 | 415 | 3.3317 | 0.1448 |
| AB2_8 | 62783 | 294 | 99.80% | 378 | 375 | 3.1403 | 0.1491 |
| BT2_1 | 43076 | 453 | 99.66% | 691 | 584 | 4.1277 | 0.0313 |
| BT2_2 | 56534 | 302 | 99.82% | 367 | 369 | 3.2740 | 0.1195 |
| BT2_3 | 62291 | 309 | 99.82% | 381 | 398 | 3.8749 | 0.0404 |
| BT2_4 | 58405 | 344 | 99.83% | 405 | 421 | 3.8326 | 0.0595 |
| BT2_5 | 53029 | 350 | 99.82% | 408 | 449 | 3.3019 | 0.1759 |
| BT2_6 | 62532 | 364 | 99.81% | 427 | 450 | 3.7203 | 0.0696 |
| BT2_7 | 61703 | 344 | 99.86% | 384 | 386 | 4.0225 | 0.0406 |
| BT2_8 | 66015 | 316 | 99.85% | 366 | 375 | 3.3736 | 0.1332 |
| CP3_1 | 59876 | 320 | 99.82% | 383 | 393 | 3.5208 | 0.0805 |
| CP3_3 | 47565 | 443 | 99.68% | 584 | 575 | 4.0316 | 0.0357 |
| CP3_4 | 50103 | 360 | 99.77% | 455 | 445 | 3.5073 | 0.1047 |
| CP3_5 | 54241 | 346 | 99.79% | 489 | 487 | 3.6179 | 0.0825 |
| CP3_6 | 48772 | 347 | 99.81% | 421 | 425 | 3.9418 | 0.0440 |
| CP3_7 | 57383 | 356 | 99.83% | 412 | 414 | 4.0763 | 0.0457 |
| AB3_1 | 59127 | 310 | 99.84% | 364 | 363 | 3.5352 | 0.0963 |
| AB3_2 | 65449 | 337 | 99.82% | 399 | 425 | 3.8225 | 0.0591 |
| AB3_3 | 49886 | 318 | 99.84% | 373 | 370 | 3.3312 | 0.1312 |
| AB3_5 | 60484 | 699 | 99.56% | 840 | 829 | 4.0077 | 0.0648 |
| AB3_6 | 52791 | 409 | 99.74% | 513 | 499 | 3.9288 | 0.0452 |
| AB3_7 | 48306 | 359 | 99.78% | 448 | 454 | 3.9420 | 0.0405 |
| BT3_1 | 47618 | 378 | 99.78% | 522 | 483 | 4.1298 | 0.0386 |
| BT3_2 | 52132 | 304 | 99.86% | 343 | 344 | 3.3575 | 0.1045 |
| BT3_3 | 54880 | 318 | 99.84% | 371 | 381 | 3.3015 | 0.1505 |
| BT3_4 | 58274 | 327 | 99.85% | 373 | 380 | 3.4893 | 0.1290 |
| BT3_5 | 51244 | 345 | 99.78% | 429 | 465 | 3.4515 | 0.0844 |
| BT3_8 | 61440 | 332 | 99.79% | 416 | 417 | 2.8887 | 0.2103 |
