## Supplementary material for "Effects of immunomodulatory peptides derived from a soil bacterium on cecal microbiota of broilers challenged with *Clostridium perfringens*": Table S2

**Table S2. List of OTUs at least 10% more enriched in each treatment at T3 sampling time**

| **OTU ID** | **Relative abundance (%)** | **Taxonomy** |
| --- | --- | --- |
| CP3 | | |
| OTU1175 | 8.66 | *Faecalibacterium* |
| OTU538 | 3.91 | *Faecalibacterium* |
| OTU1609 | 1.94 | *Ruminococcaceae_UCG-005* |
| OTU1586 | 1.37 | *Ruminiclostridium_5* |
| OTU130 | 1.31 | *Butyricicoccus pullicaecorum 1.2* |
| OTU1585 | 0.90 | *Butyricicoccus* |
| OTU687 | 0.88 | *Anaerotruncus* |
| OTU909 | 0.76 | *Ruminococcaceae_UCG-014* |
| OTU722 | 0.79 | *Ruminococcaceae* |
| OTU1161 | 0.52 | *Anaerotruncus colihominis DSM_17241* |
| OTU775 | 0.51 | *Ruminococcaceae_UCG-014* |
| OTU530 | 1.09 | *Lachnospiraceae* |
| OTU317 | 0.76 | *Tyzzerella* |
| OTU916 | 0.76 | *Lachnospiraceae* |
| OTU576 | 0.50 | *Lachnospiraceae* |
| OTU1111 | 10.68 | *Alistipes sp. CHKCI003* |
| OTU377 | 8.79 | *Barnesiella* |
| OTU1606 | 0.53 | *Clostridiales_vadinBB60_group* |
| AB3 | | |
| OTU1092 | 4.15 | *Ruminococcaceae* |
| OTU933 | 3.65 | *Ruminococcaceae* |
| OTU1549 | 1.66 | *Ruminococcaceae_UCG-008* |
| OTU1037 | 0.85 | *Butyricicoccus* |
| OTU306 | 0.82 | *Ruminiclostridium* |
| OTU797 | 0.64 | *Ruminococcaceae_UCG-004* |
| OTU678 | 2.05 | *Clostridiales_vadinBB60_group* |
| OTU1534 | 0.54 | *Clostridiales_vadinBB60_group* |
| OTU1634 | 1.72 | *Mollicutes_RF9* |
| OTU910 | 0.89 | *Bacillales* |
| OTU1603 | 0.83 | *Gastranaerophilales* |
| OTU762 | 0.80 | *Senegalimassilia* |
| OTU565 | 0.76 | *Anaeroplasma* |
| OTU1104 | 0.61 | *[Ruminococcus]_torques_group* |
| BT3 | | |
| OTU810 | 28.03 | *Bacteroides dorei* |
| OTU668 | 2.40 | *Ruminococcaceae* |
| OTU587 | 1.75 | *Ruminococcaceae* |
| OTU1637 | 0.90 | *Ruminococcaceae* |
| OTU1507 | 2.23 | *Eisenbergiella* |
| OTU568 | 0.86 | *Lachnospiraceae* |
| OTU566 | 0.64 | *[Ruminococcus]_torques_group* |
| OTU1648 | 1.41 | *Gastranaerophilales* |
| OTU1636 | 0.64 | *Gastranaerophilales* |
| OTU1186 | 3.09 | *Alistipes* |
| OTU1116 | 0.65 | *Mollicutes_RF9* |
| OTU1114 | 0.65 | *Clostridiales_vadinBB60_group* |
